# Engineering *Candida boidinii* formate dehydrogenase for activity with NMN(H)

**DOI:** 10.1101/2024.07.17.604001

**Authors:** Salomon Vainstein, Scott Banta

## Abstract

Multi-step enzymatic reaction cascades often involve cofactors that serve as electron donors/acceptors in addition to the primary substrates. The co-localization of cascades can lead to cross-talk and competition, which can be unfavorable for the production of a targeted product. Orthogonal pathways allow reactions of interest to operate independently from the metabolic reactions within a cell; non-canonical cofactor analogs have been explored as a means to create these orthogonal pathways. Here, we aimed to engineer the formate dehydrogenase from *Candid boidinii* (CbFDH) for activity with the non-canonical cofactor nicotinamide adenine mononucleotide (NMN(H)). We used PyRosetta and structural alignment to design mutations that enable CbFDH to use NMN^+^ for the oxidation of formate. Although the suggested mutations did not result in enhanced activity with NMN^+^, we found that PyRosetta was able to easily design single mutations that disrupted all enzymatic activity.

## Introduction

Orthogonal enzyme pathways have generated considerable interest due to their ability to direct specific chemical transformations within complex biological systems (Campos et al., 2019). These orthogonal pathways allow for the coexistence of distinct enzymatic cascades that operate distinctly from other metabolic pathways and reactions occurring in a cell. Orthogonal pathways achieve functional independence through various mechanisms, from compartmentalization, spatial segregation, and cofactor specificity. Compartmentalization confines reactions to specific subcellular compartments or organelles, helping avoid cross-reactions (Pei et al., 2022). Spatial segregation is achieved through protein-protein interactions or through the organization of enzymes in macromolecule complexes (Van Fossen et al., 2019). Substrate specificity mechanisms, involving differences in substrate recognition and affinity, also allow enzyme cascades to operate without interfering with surrounding reactions in a cell (Dorr et al., 2014). Orthogonal pathways have naturally evolved within cells; one example is the nicotinamide adenine dinucleotide (NAD(H)) and nicotinamide adenine dinucleotide 2’-phosphate (NADP(H)) cofactors. These two biomolecules are structurally similar, with the only difference being a phosphate group attached to the 2’ adenine ribose on the NADP(H). Both cofactors perform the same redox functions, but enzymes have evolved to differentiate the minor phosphate modification, creating natural orthogonal pathways (Cahn et al., 2017).

Oxidoreductases can have a cofactor-dependence, requiring two or more substrates for their reaction to proceed (Harel et al., 2014). These enzymes are involved in a wide variety of essential processes, including energy production, metabolism, and cellular signaling. The most common cofactors associated with oxidoreductases are NAD(P)(H) and flavin adenine dinucleotide (FAD(H2), which accept or transfer electrons with a primary substrate (Macheroux et al., 2011). Oxidoreductases have been exploited in various biotechnological processes due to their selectivity and efficiency; targeting oxidoreductases for the creation of novel orthogonal pathways would be beneficial for the production of organic products by reducing interference between competing reactions.

NAD(H) and NADP(H) are well-established canonical cofactors that play essential roles in cells, but many reactions compete for them. One method to create orthogonal pathways is to shift the cofactor specificity of the natural enzyme to a novel non-canonical cofactor that is not utilized by any other enzyme in a cell. Engineering enzymes for activity with non-canonical cofactors is being conducted by many research groups in an effort to create orthogonal reactions. There exists a database of cofactors and their functions, which can be useful in identifying potential non-canonical candidates (Richter, 2013). Many of these non-canonical cofactors need to be chemically synthesized before either being used for *in vitro* application, or fed into living cells (Weusthuis et al., 2020). Designing biological synthesis motifs to allow for non-canonical cofactors to accumulate within cells is essential for reducing chemical synthesis costs and to circumvent the challenges delivering molecules through a cell membrane. For example, the Zhao group showed that they could accumulate nicotinamide cytosine dinucleotide (NCD(H)), an NAD(H) analog, in cells by engineering and overexpressing an NCD synthetase (Wang et al., 2021). They then showed that the NCD(H) cofactor was utilized as a cofactor *in vivo* to convert L-malate to D-lactate.

Recent studies have shed light on the emergence of another intriguing molecule, nicotinamide mononucleotide (NMN(H)), as an *in vivo* non-canonical cofactor source. NMN^+^ is an intermediate molecule in the biosynthesis of NAD^+^. A visualization of NAD^+^ sectioned into its components is shown in Figure 1. Although most enzymes do not use NMN(H) as a cofactor, recent studies have shown that NMN(H) can act as a signaling molecule by directly activating sirtuins, which regulate metabolism, aging and stress response (Tuncay et al., 2023, Imai, 2010). NMN(H) has been considered as a therapeutic in the hopes of improving NAD^+^ synthesis in age related diseases and neurogenerative disorders. Although NMN(H) is present within cells and serves as a signaling molecule, there is little evidence of its use as a cofactor in oxidoreductases reactions; therefore, NMN(H) is a prime candidate to serve as orthogonal cofactor. The Banta lab engineered an alcohol dehydrogenase to use (NMN(H)), which proved useful in an enzymatic biofuel cell application (Campbell et al., 2012). The Li lab has worked with cell lines with NAD^+^ synthesis knocked out, leading to NMN^+^ accumulation within a cell (Black et al., 2020a, Black et al., 2020b). They then used computational and rational approaches to design a glucose dehydrogenase (GdH) that was able to use the NMN^+^ as a cofactor in an essential metabolic cascade. The mutations revealed a general design strategy for enzymes containing the Rossmann fold cofactor-binding motif of the GdH where the mutations mimic the steric interactions that would be present with an NAD(P)^+^ cofactor. The Li lab has also designed a glutathione reductase which specifically uses NMNH, and used directed evolution approaches to engineer a phosphite dehydrogenase to reduce NMN^+^ (Zhang et al., 2022).

**Figure 1.**
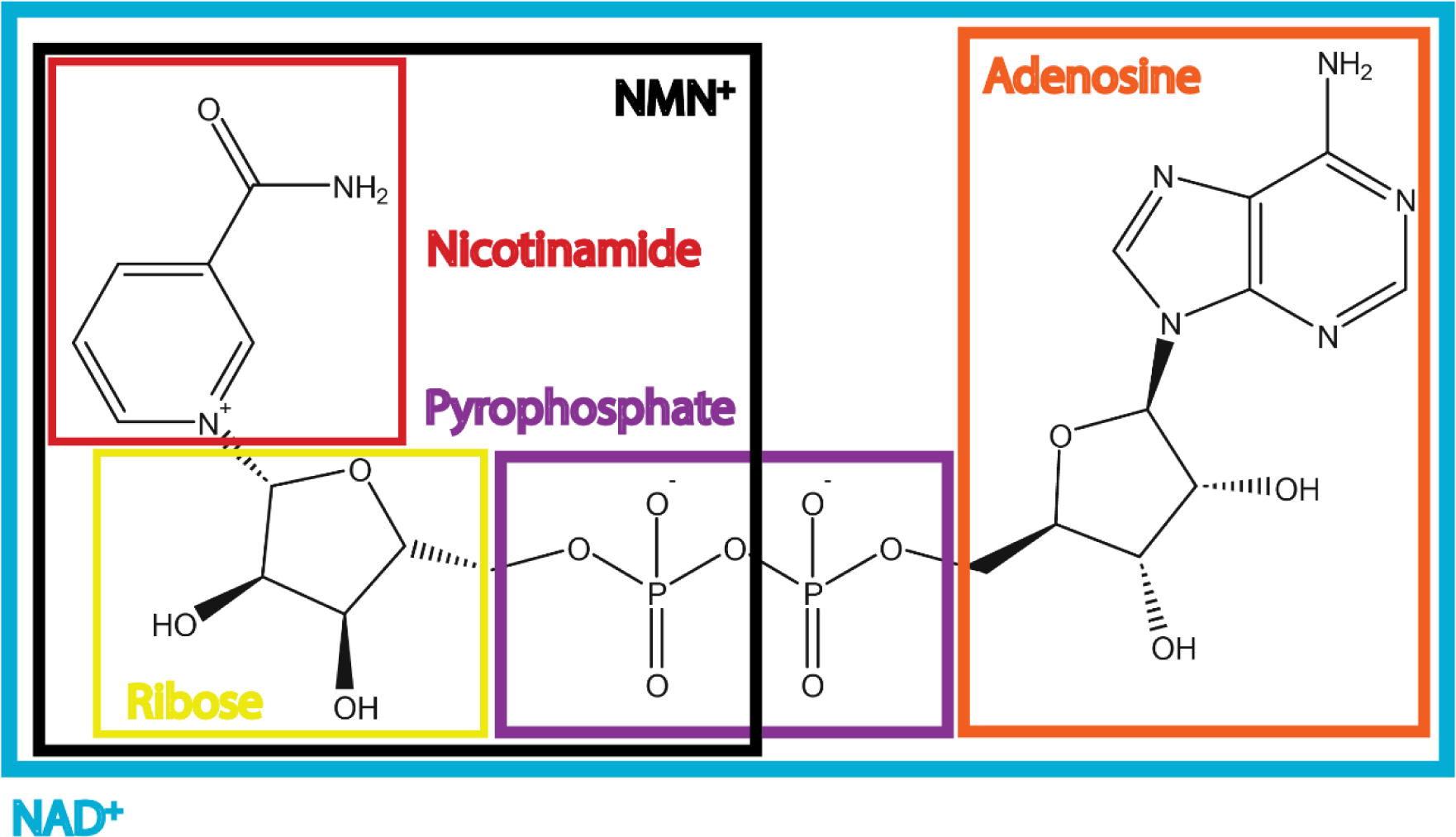
Structure of NAD+ with key sections highlighted. The full molecule, outlined in light blue, is an oxidized NAD+. The nicotinamide is boxed in red, the nicotinamide ribose is boxed in yellow, the pyrophosphate is boxed in purple, and the adenosine (comprised of a ribose and adenine) are boxed in orange. NMN+ contains the nicotinamide, ribose, and one phosphate group as shown in the black box. The fourth oxygen on the phosphate group would carry a negative charge.

Most enzymes do not have natural affinity or activity with non-canonical cofactors, including analogs which are structurally and chemically similar to the original cofactor. Research groups have been working on altering cofactor specificity for decades, with varying degrees of success. The Baker lab has worked on designing enzymes *de novo* for activity with specific and novel substrate (Donohoe et al., 2013). However, in most cases, an existing enzyme is used as a starting point, and mutations are made at different positions to guide the enzyme to not only bind to a specific cofactor, but to be able to use it for the reaction. These mutations are determined in a number of ways, but are generally broken down into a directed evolution approach, rational design, or some hybrid of the two (Brannigan and Wilkinson, 2002, Bilal et al., 2018, Rajeshwari and Pratyoosh, 2019). Rational design is attractive; however, it may not always identify useful mutations. For example, when engineering the cofactor selectivity of an enzyme between NAD^+^ and NADP^+^, it may seem like mutations around the 2’-phosphate group would offer the best approach, but it has been shown that distal mutations can also lead to improved selectivity (Solanki et al., 2016, Solanki et al., 2017). It is difficult to account for every interaction among the amino acids in a protein, the protein interactions with the substrate, and protein interactions with the surrounding environment. Some computational programs rely on a structure-based strategy for designing mutations and altering protein dynamics, while other programs are more data driven and rely on machine learning algorithms to predict functional mutations (Mazurenko et al., 2019). PyRosetta combines a Python-based interface to the Rosetta molecular modeling platform; it has the ability to estimate protein energies, perform docking simulations, explore effects of mutations, and find lowest energy conformations (Chaudhury et al., 2010, Le et al., 2021).

In this work, we attempted to use PyRosetta to engineer the formate dehydrogenase from *Candida boidinii* (CbFDH) to accept NMN(H) as a non-canonical cofactor. Formate is an inexpensive reductant and the CbFDH dehydrogenase has been used in a number of cofactor regeneration processes (Hoelsch et al., 2013). Like the GdH enzyme, CbFDH contains a Rossmann fold, a super secondary structure that is found across many oxidoredcutases (Hanukoglu, 2015). The successful mutations made in the GdH enzyme (Black et al., 2020b) were used as a guide for designing CbFDH for activity with NMN(H). The efficacy of this computational design approach is discussed.

## Materials and Methods

### Materials

The primers used for mutations were synthesized by IDT DNA. BL21(DE3) and 5-alpha competent *E. coli* were purchased from New England Biolabs (NEB). Q5 site-directed mutagenesis kit was from NEB. 96-well UV plates used in the assays were bought from Thermo Fisher. Absorbance and fluorescence measurement were performed on a SpectraMax iD5 plate reader from Molecular Devices. The QuickDrop micro-volume spectrophotometer was from Molecular Devices. DNA extraction was performed using the spin miniprep kit from Qiagen. Formate, cofactors, and other chemicals were purchased from Sigma.

### Rational and Computational Design

The crystal structure for CbFDH in complex with NAD^+^ and azide (PDB: 5DN9) was visualized using PyMOL. Although the enzyme exists as a homodimer, the monomer is shown in Figure 2 to better visualize the active site. Computational modelling and mutation simulations were performed using PyRosetta3. An algorithm was developed that allows user input of residues for mutation, and PyRosetta was used to perform saturation single mutagenesis at each of these positions, and to find the minimum energy conformation with a repacking radius of 8 Å (Vainstein and Banta, 2023). One algorithm was designed to prioritize improving the specificity for NMN^+^, and the output was a list improved mutants based on the ratio of the energy score with NMN^+^ versus NAD^+^. A second algorithm prioritized the energy score with NMN^+^ and did not take into account a specificity score. A params file was constructed for NMN^+^ for use in PyRosetta for docking and mutation simulations. The NAD params file was extracted, and the adenosine end half was removed, resulting in an NMN^+^ molecule with the same conformation and localization as the original nicotinamide from NAD^+^; leaving the NMN^+^ at the same location was important to design mutations around it that would allow the enzyme to retain its original catalytic function. The CbFDH with docked NAD^+^ and NMN^+^ can be seen in Figure 2a and Figure 2b respectively. The updated PDB file with a docked NMN^+^ was loaded into PyRosetta and repacked to obtain the lowest energy conformation. The sequence section associated with the PDB entry was used to identify residues comprising the NAD(H) binding site and the formate binding site (Figure 2c) which determined their inclusion in the mutation program; the orange residues are on the nicotinamide side of the cofactor, and were not used in order to avoid interfering with substrate interactions. Residues shown in grey are along the pyrophosphate and residues shown in blue surround the adenosine; both of these groups were considered for design.

**Figure 2.**
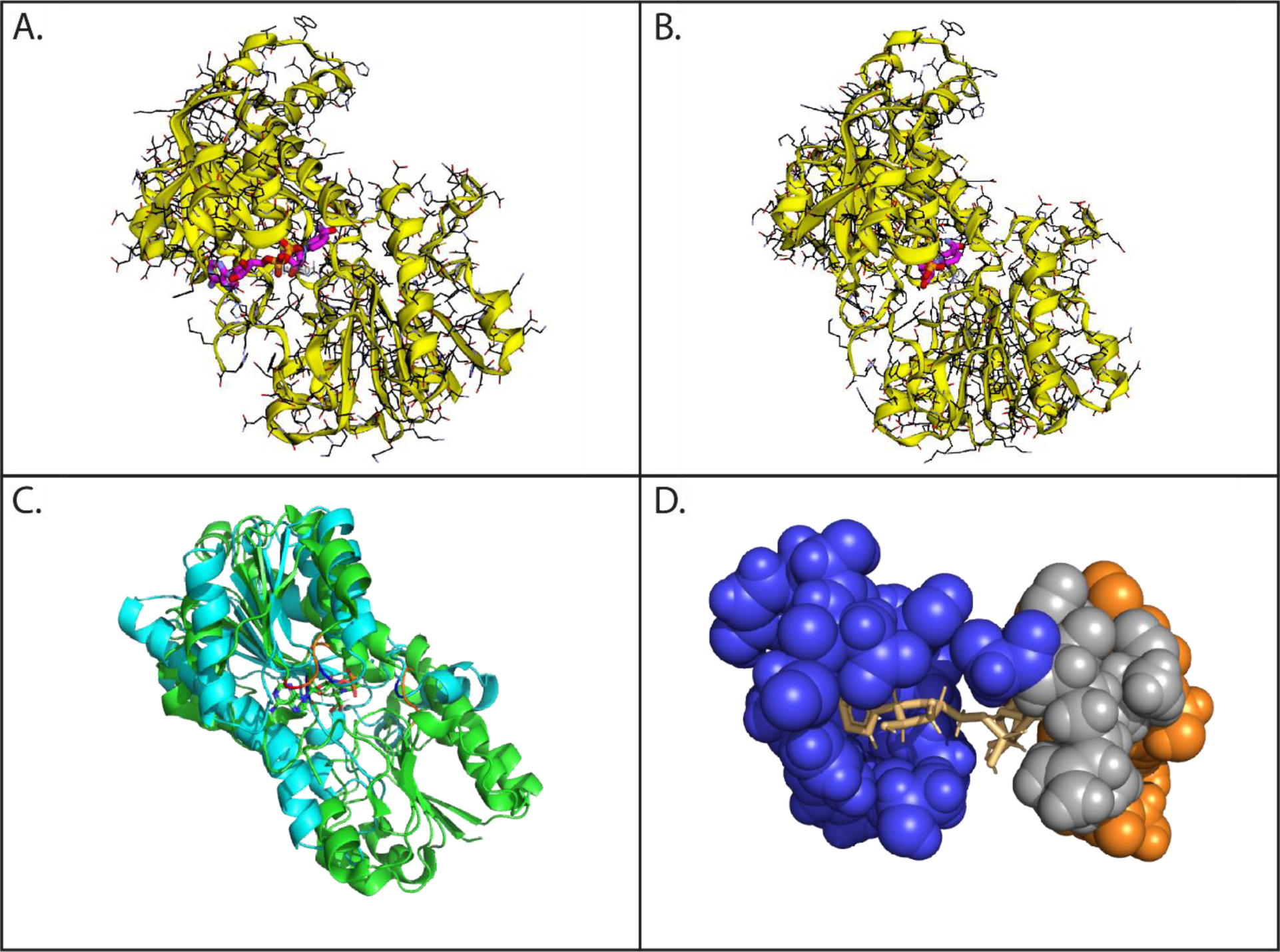
A) The crystal structure of a single monomer of CbFDH with the docked NAD+ (PDB: 5DN9), and visualized using PyRosetta. B) The crystal structure of a single monomer of CbFDH with the docked NMN+, and visualized using PyRosetta. For panels A and B, the enzyme is shown in yellow, and the cofactor is shown in pink, orange and red. C) Alignment of CbFDH (green) with GdH (cyan) from Black et al., with a bound NAD+ shown with element fill. D) A closer view of the binding pocket of CbFDH with neighboring residues shown that interact with the cofactor; orange residues (P68, P69, G92, N119, V123, D282, V283, H311, S313, G314) surround the nicotinamide ring, grey residues (V93, V120, R174, I175) surround the pyrophosphate and blue residues (G171, G173, Y194, D195, Y196, A229, P230, H232, T235, A357) surround the adenosine.

CbFDH was structurally aligned with the GdH from Black et al. (Figure 2d). Both enzymes contain a Rossmann fold motif and the successful mutation made on the GdH was used to offer insight to demine other positions that may be important for mutations.

### Cloning and transformation of mutant genes

The CbFDH gene (Willett et al., 2022) was cloned into the pET28a vector with a His-Tag on the N-terminus for immobilized metal affinity purification. The CbFDH-pET28a plasmid was transformed into BL21(DE3) *E. coli.* Mutation primers were designed using NEBuilder to create the best scoring mutations provided by the PyRosetta analysis. All primers used are listed in the SI (Table S1). Mutations were incorporated via PCR following the Q5 site-directed mutagenesis protocol. The PCR product was added to the KLD enzyme kit following the included protocol. The resulting DNA product was transformed into 5-alpha competent *E. coli* cells following the provided protocol. The cells were plated on Luria broth (LB) agar plates containing 50 μg/mL of kanamycin sulfate and incubated overnight at 37 °C. Colonies from overnight plates were picked and inoculated into 5 mL of LB (50 μg/mL kanamycin sulfate) and were incubated (37 °C, 250 rpm) overnight.

The overnight cultures were miniprepped following the QIAprep spin miniprep protocol. DNA concentrations were quantified via nanodrop and DNA samples were sequenced and compared to the wild type using the basic local alignment search tool (BLAST). Sequences that best incorporated the correct mutations were transformed into BL21(DE3) *E. coli* following the NEB protocol, and plated on LB agar (50 μg/mL kanamycin). Plates were incubated at 37 °C overnight. Single colonies were picked from the plate and grown in 5 mL LB (50 μg/mL kanamycin) overnight. The following day, 0.5 mL of saturated overnight was cryopreserved (-80 °C).

### Expression and purification for initial screening

Overnight cultures were used to inoculate 50 mL of Terrific Broth (TB, 50 μg/mL kanamycin sulfate) in a 250 mL Erlenmeyer flask, which was placed in an orbital shaking incubator at 37 °C until an optical density at 600 nm (OD600) of ∼0.6 was reached. Protein expression was induced by adding isopropylthio-β-galactoside (IPTG) to a final concentration of 1 mM. The cultures were expressed at 30 °C overnight.

The following morning, cultures were centrifuged (4000 xg, 12 minutes, 4 °C) and the supernatants were discarded. Pellets were resuspended in 4 mL of Wash Buffer A (20 mM Tris, 300 mM NaCl, pH 7.4), then divided equally into microcentrifuge tubes (1.5 mL volume) and spun at 17,000 xg for 3 minutes at 4 °C. Supernatants were discarded, and the pellets were stored at -20 °C.

Thawed cell pellets were lysed with 800 μL of Cell Lytic B Cell Lysis Reagent, and the samples were rotated at room temperature for 20 minutes. Tubes were centrifuged at 17,000 xg for 30 minutes at 4 °C. The protein was recovered from each lysate using HisPur Ni-NTA Magnetic Beads (ThermoFisher). Briefly, 160 μL of beads were pipetted into 1.5 mL Eppendorf tubes, which were then placed on a magnetic rack. The supernatant (storage solution) was removed and the beads were washed with 800 μL of Wash Buffer A. 400 μL of Wash Buffer A and 400 μL of enzyme lysate were combined to create a half-diluted solution of the lysate in wash buffer to the magnetic beads. The tubes were vortexed for ten seconds, and then placed on a rotator at 4 °C for one hour. Tubes were placed on the magnetic rack, and the supernatant was removed. Beads were washed with 800 μL of Wash Buffer B (20 mM Tris, 300 mM NaCl, 40 mM imidazole, pH 7.4) with a ten second vortex, placed on the magnetic rack, and the supernatant was removed. To elute the protein, 50 μL of Elution Buffer (20 mM Tris, 300 mM NaCl, 500 mM imidazole, pH 7.4) was added, and the tubes were vortexed for ten seconds every five minutes over a period of 15 minutes. The samples were placed in the magnetic rack, and the supernatant was removed and saved in a separate tube. This elution procedure was repeated a second time, ending with a final volume of 100 μL of eluted protein. Protein concentrations were measured with a nanodrop using an extinction coefficient of 51,402 M^-1^ cm^-1^. Magnetic beads were washed and stored in ethanol (20% v/v).

### Activity assays and full kinetic experiments

Initial screening assays were conducted to identify mutants with activity with NMN^+^, as well as NAD^+^ and NADP^+^. A cofactor reaction mix was made for each cofactor containing 600 μM of cofactor and 40 mM sodium formate in 50 mM Tris Buffer (pH 7.2). Purified enzymes were diluted to 500 nM in 50 mM Tris Buffer (pH 7.2). 100 μL of the enzyme reaction mix was pipetted into a 96 well UV plate and 100 μL of the enzyme mix was added to initiate the reactions. The final reaction conditions were 300 μM cofactor, 20 mM sodium formate, and 250 nM enzyme in 50 mM Tris Buffer (pH 7.2), measured at 25 °C. Absorbance at 340 nm was followed over time. Screening measurements were conducted in duplicate.

## Results

### CbFDH mutant designs

PyRosetta was used to explore site-directed mutations in the CbFDH sequence at positions proximal adenosine and pyrophosphate groups of the docked NMN^+^ cofactor. The best scoring mutants were Y196M, Y196I, and Y196L. When considering a specificity shift from NAD^+^ to NMN^+^, the best scoring mutants were D195W, G173I, and G173L with specificity scores of -11, -6, and 29 respectively. A negative specificity scores suggest that the mutation was very unfavorable for enzyme affinity with the NAD^+^ cofactor, leading to a positive PyRosetta energy value for the NAD^+^-CbFDH complex, while the score for the NMN^+^-CbFDH complex remained negative. Mutants were expressed and assayed for activity with NAD^+^, NADP^+^, and NMN^+^.

Structural alignment with the previously engineered GdH offered additional suggestions for mutations that would lead to NMN^+^ activity. These were: V93R, T116R, G117R, Y196Q, A229K, and Y358Q.

### Activity Screening

Absorbance measurements were taken for the mutants and three cofactors over time. Initial rates were calculated as the slope of the linear region of the duplicate data, with errors calculated using Excel (Figure 3). Any negative values for activity were recorded as zero. The wildtype enzyme (WT) with formate and NAD^+^ had the highest specific reaction rate. Mutants H232Y and Y196M had a ∼20% decrease in initial specific rate with NAD^+^ compared to the WT. Surprisingly, many mutations had no apparent activity with NAD^+^, with D195W being inactive with NAD^+^. The highest initial rates with NMN^+^ were two orders of magnitude lower than those with NAD^+^, with H232N having the largest value of 0.0038 s^-1^. The largest activity with NADP^+^ was also H232N, with a value of 0.0072 s^-1^. These values are dramatically lower compared to the original WT activity. Overall, none of the mutants showcased an increased level of activity with NMN^+^ under the conditions tested.

**Figure 3.**
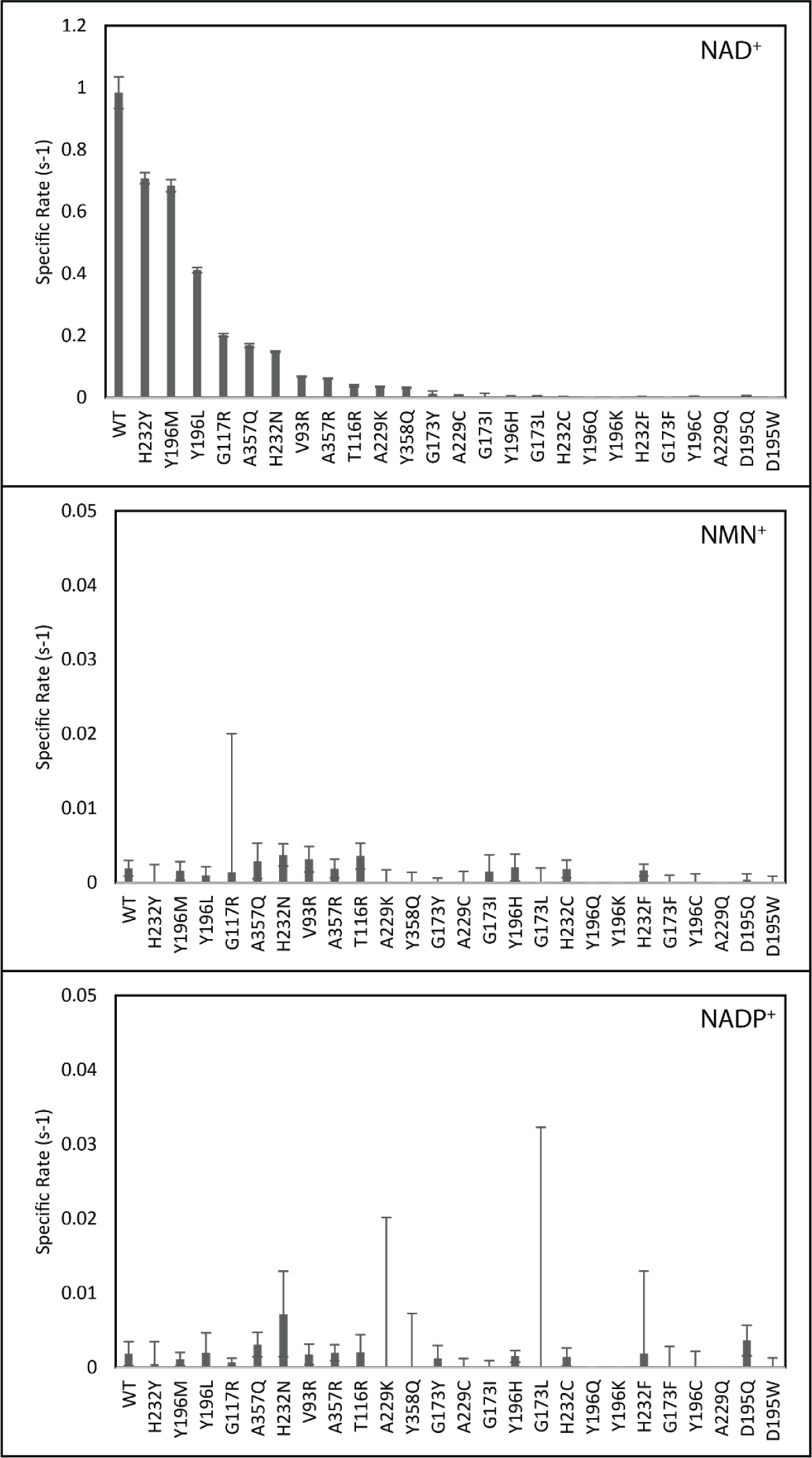
Initial specific rates of the single mutants. Reaction conditions included 300 μM cofactor, 20 mM sodium formate, and 250 nM enzyme in 50 mM Tris Buffer (pH 7.2), measured at 25 °C. Data were measured in duplicate, and error bars represent the standard error of the linear regression. The y-axis is scaled to 1.2 s^-1^ for NAD^+^ as a cofactor, and 0.05 s^-1^ for NMN^+^ and NADP^+^ as the cofactor. The largest specific rate at these conditions was achieved by the WT enzyme utilizing NAD^+^.

## Discussion

Altering the cofactor specificity of an enzyme for a non-canonical cofactor can enable the creation of orthogonal reaction pathways *in vivo*. Non-canonical cofactors have traditionally been incorporated into cells by feeding the cofactors to growing cultures (Zachos et al., 2019). However more recently, there have been efforts aimed at engineering cell lines that can produce non-canonical cofactors. Engineering enzymatic pathways to specifically use the non-canonical cofactors allows for the creation of novel orthogonal cascades with exciting biotechnology applications.

Theoretical approaches to protein engineering have been around for decades and began as free energy calculations to predict how mutations would alter the thermodynamics within a protein and with protein interactions (Marabotti et al., 2021). Computation approaches to protein design continue to be a growing trend with varying degrees of success, especially when factoring in non-proteogenic compounds such as cofactors, post-translational modifications, and nucleic acids (Korbeld and Furst, 2023). Algorithms have been created to predict enzyme structures from the amino acid sequences, predict binding sites, dock substrates, remodel the active sites, and generate potentially useful mutant libraries. However, limited computational power and relatively little available data for machine learning algorithms leads to questionable reliability in any computational result (Korbeld and Furst, 2023).

We focused on engineering CbFDH for activity with NMN(H), a precursor to NAD^+^ that is present within cells, but does not serve as an enzymatic cofactor. The presence of NMN(H) is benign to endogenous enzymes, allowing essential metabolic processes to continue uninterrupted. Mutations were determined using two main approaches: structural alignment with a previously engineered enzyme and through predictions made using PyRosetta. Black et al. previously designed GdH for activity with NMN(H) (Black et al., 2020b), and this enzyme is an oxidoreductase which contains the Rossmann fold found in many other enzymes including CbFDH. Structural alignment was performed to match the successful mutations on GdH to similar positions on CbFDH in an attempt to discover a general design principle that would result in NMN^+^ activity gain. However, none of these mutations led to improved functional activity with the non-canonical cofactor NMM^+^.

PyRosetta was used to explore two avenues for mutant discovery, one that was focused on changing selectivity and one that was focused on improving affinity. The mutants suggested for shifting selectivity to NMN^+^ did not lead to functional activity with NMN^+^, but instead reduced the activity with NAD^+^. The D195W was the mutant with the largest predicted shift in selectivity, however this mutation eliminated activity with NMN^+^, NAD^+^ and NADP^+^ (Figure 3).

Engineering proteins for activity with novel cofactors is a complex problem and, the computational scores provided by PyRosetta did not result in enhance activity levels. The changes in the energy score should correlate to the binding or affinity to the cofactor; however, high affinities may lead to slow cofactor release, inhibiting turnover. A comparison of the PyRosetta scores and the experimental specific activities is shown in Figure 2.4. There is no correlation between the PyRosetta energy score and the resulting activity that was measured.

**Figure 4.**
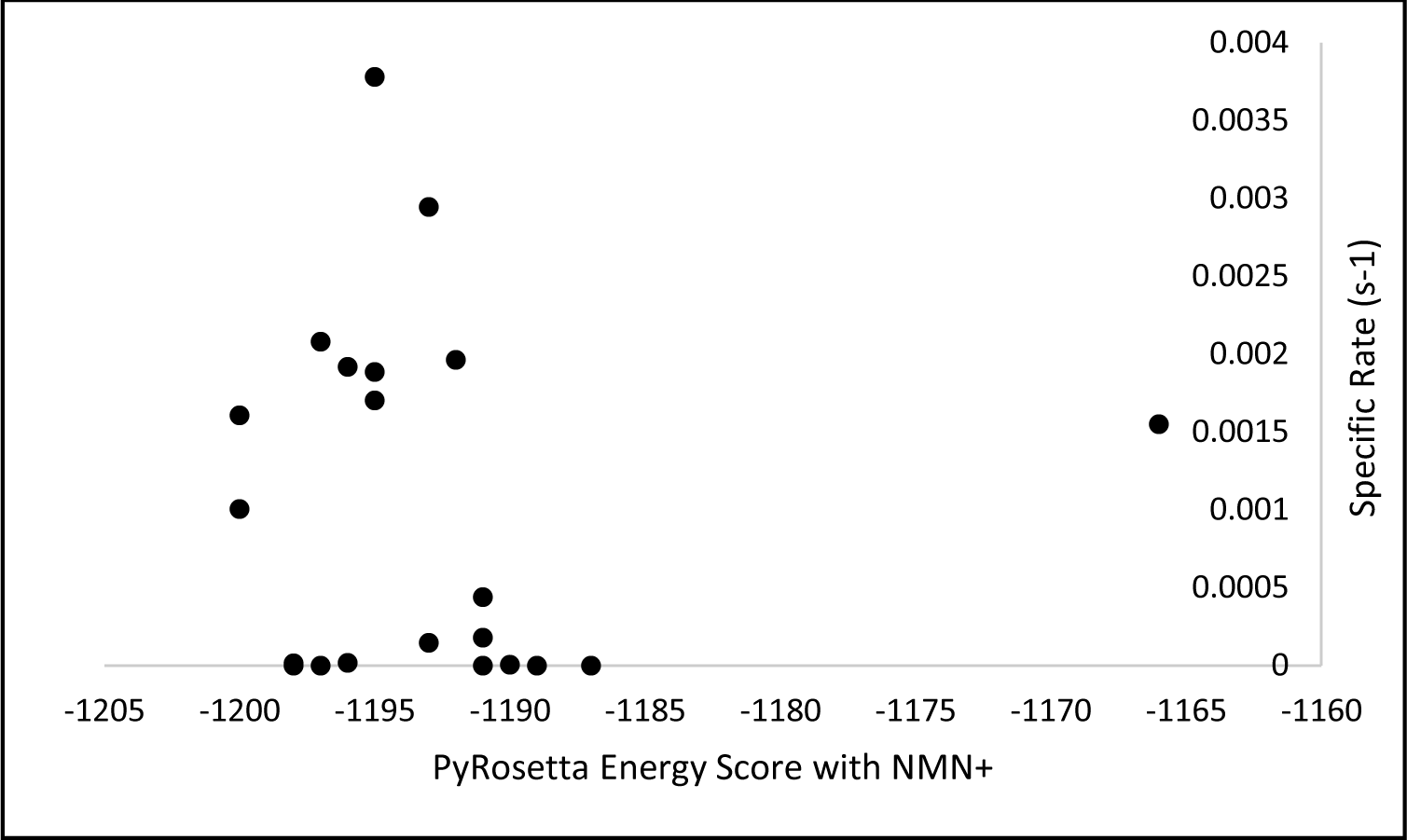
Specific rates measured experimentally versus PyRosetta energy scores.

In most cases, mutating a single amino acid in a protein leads to minor changes, and it was interesting that many mutations suggested by the PyRosetta algorithm led to dramatic decreases in activity for NAD^+^. The PyRosetta algorithm only explored residues involved with NAD^+^ binding, some of which are conserved residues among Rossmann fold enzymes; this could explain why a single mutation could have devastating consequences to enzymatic activity. The Rossmann fold is characterized by glycine rich loop domains between the alpha helices and beta sheets (Bossemeyer, 1994), so mutations such as G173 may destabilize the NAD^+^ binding site. The PyRosetta energy score output values did not result a large variability among the top scoring mutants, however the score increased and reached positive values for some of the worst performing mutants, suggesting that a strength of this computational approach may be in locating positions that are vital for enzymatic activity and for recommending mutations for activity disruption.

## Supplementary Information

**Table S1.**
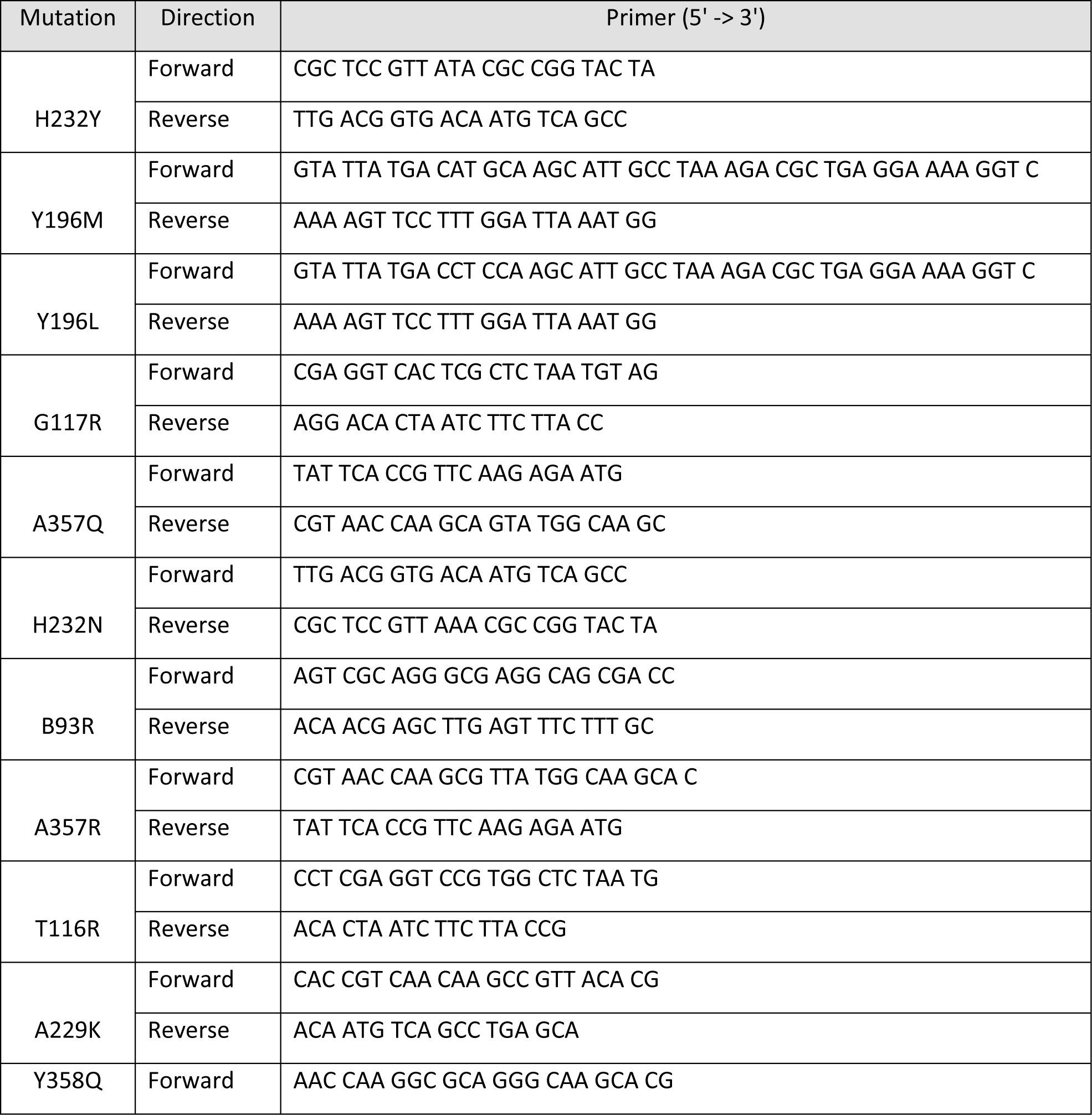
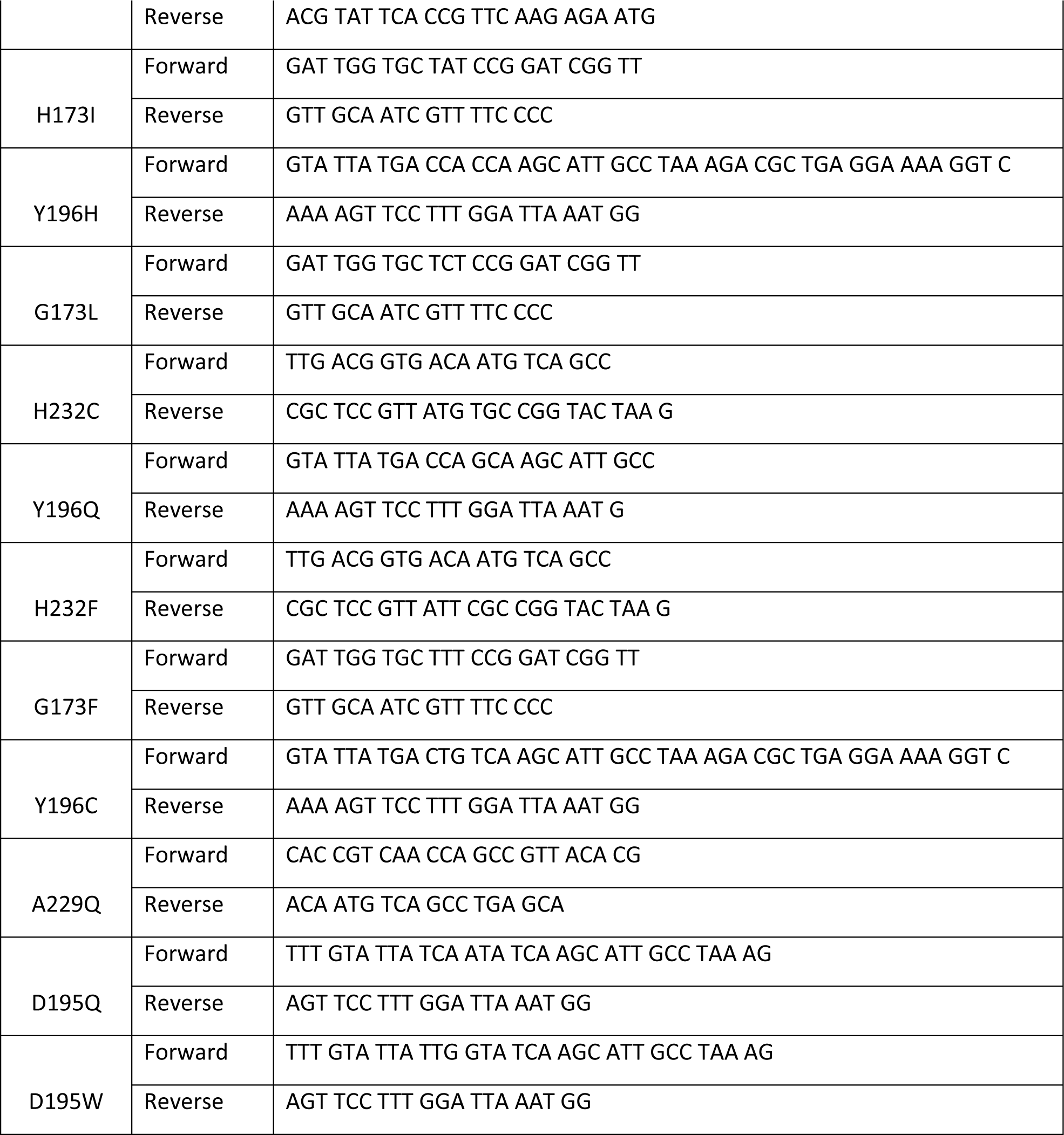
Mutation primers for the single mutants made to assess activity with NMN^+^. Forward and reverse primers are shown, and both are listed in the 5’ to 3’ direction.

